# A formin-mediated cell wall- plasma membrane- cytoskeleton continuum is required for symbiotic infections in *Medicago truncatula*

**DOI:** 10.1101/2020.07.10.197160

**Authors:** Pengbo Liang, Clara Schmitz, Beatrice Lace, Franck Anicet Ditengou, Jean Keller, Cyril Libourel, Pierre-Marc Delaux, Thomas Ott

## Abstract

Plant cell infections are tightly orchestrated by cell wall (CW) alterations, plasma membrane (PM) resident signalling processes and dynamic remodelling of the cytoskeleton. During root nodule symbiosis these processes result in morpho-dynamic responses including root hair swelling and curling, PM invagination and polar growth of a tubular infection structure, the infection thread (IT). However, the molecular details driving and guiding these PM remodelling events remain to be unravelled. Here, we studied a formin protein (SYFO1) in *M. truncatula* that is specifically induced during rhizobial infection. Phenotypical analysis of *syfo1* mutants clearly indicates that the encoded protein is required for efficient rhizobial colonization of root hairs. SYFO1 itself creates a proteinaceous bridge between the CW and the polarized cytoskeleton. It binds to CW components via a proline-rich N-terminal segment, which is indispensable for its function. On the cytoplasmic side of the PM SYFO1 is associated with actin accumulations supporting the hypothesis that it contributes to cell polarization *in vivo*. This is further sustained by the fact that cell shape changes can be induced in a stimulus-dependent manner in root protoplasts expressing SYFO1. Taken together we provide evidence for the evolutionary re-wiring of a generic cytoskeleton modulator into a symbiosis-specific response.

## INTRODUCTION

Legumes have the unique ability to symbiotically associate with rhizobia to maintain a nitrogen-fixing mutualism. Intracellular colonization of *Medicago truncatula* roots by the compatible rhizobium *Sinorhizobium meliloti* initiates from young, growing root hairs. In the course of the interaction an organogenetic program is executed in the root cortex and the pericycle that results in the development of nodules, in which symbiotic nitrogen fixation takes place [1, 2]. The first morphological step to establish this symbiosis comprises a rhizobial trap, where a growing root hair engulfs the symbiont by physically curling around it [3, 4]. Phenomena such as root hair deformation and root hair branching, steps that precede bacterial trapping in legumes [5, 6], have been generally observed in plants in response to the microtubule-stabilizing agent taxol [7], in mutants like the kinesin *mrh2* [8] or upon over-expression of the formin protein AtFH8 [9] and the ROP GTPase RHO-OF PLANTS 2 [10]. In contrast, incomplete curls were only observed at low frequency in *Arabidopsis* mutants affected in the loci *CEN1*, *CEN2*, *CEN3* [11] and in the *scn1-1* mutant, where the corresponding locus encodes the RhoGTPase GDP dissociation inhibitor SUPERCENTIPEDE1 [12]. This indicates full root hair curling to represent a rather specific invention. The entrapment of the symbiont completes with its full enclosure between root hair cell walls in a structure called the ‘infection chamber’ (IC) [3–5, 13]. This is followed by exocytotic secretion of host-derived cell wall loosening enzymes such as NODULE PECTATE LYASE [14] which, most likely, subsequently allows the formation of a negatively curved plasma membrane (PM) structure that further elongates into a tube-like channel, the ‘infection thread’ (IT) [3, 4, 15]. Parallel to these morphological changes a set of transcription factors genetically re-programs host root cells to allow transcellular IT progression and nodule organogenesis [16–22].

Molecularly, initial root hair responses are triggered upon the recognition of bacterial signalling molecules, called Nod Factors, by host LysM-type receptor-like kinases [23–28]. These include root hair swelling, deformation and branching that precede root hair curling. This is aided by a tip-localized cytosolic calcium gradient [29, 30], global actin re-arrangements and dense subapical fine actin bundles that are required for the delivery of Golgi-derived vesicles to the root hair tip [5, 31–33]. However, the molecular machinery steering actin reorganisation and polarity during root hair curling remains elusive. Altered actin dynamics during early responses to NFs or rhizobia have been shown in mutants such *Lotus japonicus pir1/nap1* and *scarn* that are affected in the actin-related SCAR/WAVE complex [34, 35]. In addition, the presence of highly dynamic F-actin plus ends in swelling root hairs and during root hair deformation implies that newly accumulated actin physically pushes towards the PM to re-initiate the growth [36]. But, different to metazoan cells, where membrane protrusions such as filopodia are driven, among others, by formin-mediated polar growth of actin filaments [37], any membrane protrusions in plant cells ultimately require local cell wall modifications and thus this membrane-cell wall continuum to be coordinatively regulated.

Here, we identified SYMBIOTIC FORMIN 1 (SYFO1) as an essential gene controlling symbiotic responses in root hairs. SYFO1 transiently re-localizes to the root hair tip upon inoculation of *M. truncatula* root hairs with *S. meliloti* and mediates the formation of polar actin accumulations. Its function strictly depends on the protein extracellular domain that mediates cell wall attachment. Taken together we demonstrate SYFO1 as the first player regulating the continuum between the plasma membrane and the cell wall during the onset of rhizobial infections.

## RESULTS

### Evolutionary and transcriptional patterns identify symbiotically regulated formins

Since formins are well known proteins conferring polar growth of actin filaments, we searched for candidates within this family that could control symbiotically induced root hair responses. Based on the presence of a conserved Formin Homology 2 (FH2) domain, we identified 19 candidates in the *Medicago truncatula* genome (Table S1). Using publicly available transcriptome data, two of them (*Medtr5g036540.1* and *Medtr8g062830.1*) were found to be transcriptionally up-regulated during early stages of symbiotic interactions [38]. We independently verified these data using quantitative RT-PCR (qRT-PCR) with *Medtr5g036540.1* being induced by about 60-fold at one day post inoculation (dpi) of roots with *S. meliloti* while only a weak induction could be confirmed for *Medtr8g062830.1* at 5 dpi (Fig. S1A). Therefore, we named the genes *SYMBIOTIC FORMIN 1* (*SYFO1* (*Medtr5g036540.1*, *MtrunA17_Chr5g0414941* in the v5r1.6 *M. truncatula* genome version) and SYFO1-like (*SYFO1L, Medtr8g062830.1, MtrunA17_Chr8g0364331*). Both encoded proteins contain a predicted signal peptide (SP) in the extracellular domain (ECD) followed by a single-span transmembrane domain (TMD), and FH2 domain in the cytoplasmic side.

To spatially resolve the observed transcriptional patterns for *SYFO1*, we generated a fluorescent reporter where a nuclear localized tandem GFP was driven by the endogenous *SYFO1* promoter (*ProSYFO1-NLS-2xGFP*). Consistent with the qRT-PCR results (Fig. S1A), the *SYFO1* promoter was activated at 1 dpi in root hairs and cortical cells (Fig.S1 B-C) while no activity was observed in the absence of the symbiont.

To supplement the identification strategy and to search for putative genetic redundancy, we studied evolutionary patterns within the formin family. The *SYFO1/1L* clade contains genes from all Eudicots species included in the analysis, and *SYFO1* and *SYFO1L* derived from the Papilionoideae duplication. In *A. thaliana*, three co-orthologs of *SYFO1* and *SYFO1L* are found: *AtFH4*, *AtFH7*, *AtFH8* likely deriving from the Brassicaceae triplication (Fig. S2). While AtFH4 and AtFH8 resemble a similar protein domain structure compared to SYFO1 and SYFO1L, AtFH7 has lost the signal peptide and displays an altered TMD domain. Mining the gene expression atlas of other Papillionoideae, we found *SYFO1* (in common bean, Phvul.002G077100.2) or *SYFO1L* (in *Lotus japonicus Lj0g3v0115049.1*) upregulated during nodulation, pinpointing for a potential shared function in nodulation within this clade.

Based on the phylogeny, we tested for relaxed or intensified selective pressure acting on different branches of interest in eudicot sequences (represented by circles) (Fig. S2; Supplemental Table S3). We identified relaxed selective pressure indicated by a higher overall ratio of non-synonymous (dN) vs. synonymous (dS) mutations (dN/dS) on the branch supporting the clade that includes all species that develop the root nodule symbiosis (NFN; K=0.54, LRT=85.3 and p-val < 1e-16). This relaxation was individually confirmed for Fabales (which includes legumes; K=0.66, LRT=18.3 and p-val< 14.1e-09), Cucurbitales (K=0.78, LRT=25.0 and p-val=5.6e-07) and Fagales (K=0.65, LRT=18.3 and p-val< 1.9e-05) but not for Rosales (K=1.17, LRT=7.4 and p-val< 6.7e-03) for which a slight intensification of selective pressure (lower overall dN/dS ratio) was detected. In contrast, Brassicaceae species are under strong intensification of the selective pressure (K=4.6, LRT=84.5 and p-val < 1e-16) that is not even relaxed by the triplication of the branch in this family. The relaxed selection pressure detected at the base of the NFN clade and at the base of the Fabales may both reflect neofunctionalization that would have occurred for the recruitment of SYFO1/1L in root nodule symbiosis. Interestingly, the corresponding orthologs of *Parasponia andersonii*, a non-legume tree that forms a Frankia-type symbiosis with rhizobia, are not induced during the symbiotic interaction. This further supports a possible functional specialisation of these formins in legumes /Papilionoidae [39].

### SYFO1 controls rhizobial infection and root hair responses

In order to genetically assess the function of SYFO1 and SYFO1L, we identified two independent *Tnt1* transposon insertion lines for *SYFO1* [*syfo1-1* (NF9730) at 485bp and *syfo1-2* (NF9495) at 1834bp downstream of the start codon] and *SYFO1L* [*syfo1L-1* (NF20350) at 1279bp and *syfo1L-2* (NF15608) at 1370bp downstream of the start codon] in Medicago wild type R108 (Fig. 1A). Endogenous *SYFO1* and *SYFO1L* transcripts were significantly reduced in both lines, respectively (Fig. S3), while only the *syfo1-1* and *syfo1-2* alleles showed a significant nodulation phenotype developing fewer nodules per root at 3 weeks post inoculation (wpi) with about half of them being aborted and/or white, indicative of non-functional nodules (Fig. 1B-D). In contrast, bacterial infection patterns of Medicago wild-type and *syfo1L-1* and *syfo1L-2* mutant nodules were identical (Fig. 1C, D and Fig. S4A, B). The infection zones were found to be reduced in white but elongated (Fig. S4C) or spherical (Fig. S4D) *syfo1* mutant nodules. The latter failed to maintain a persistent meristem (Fig. S4D). Therefore, we selected SYFO1 as the prime candidate of interest.

**Figure 1.**
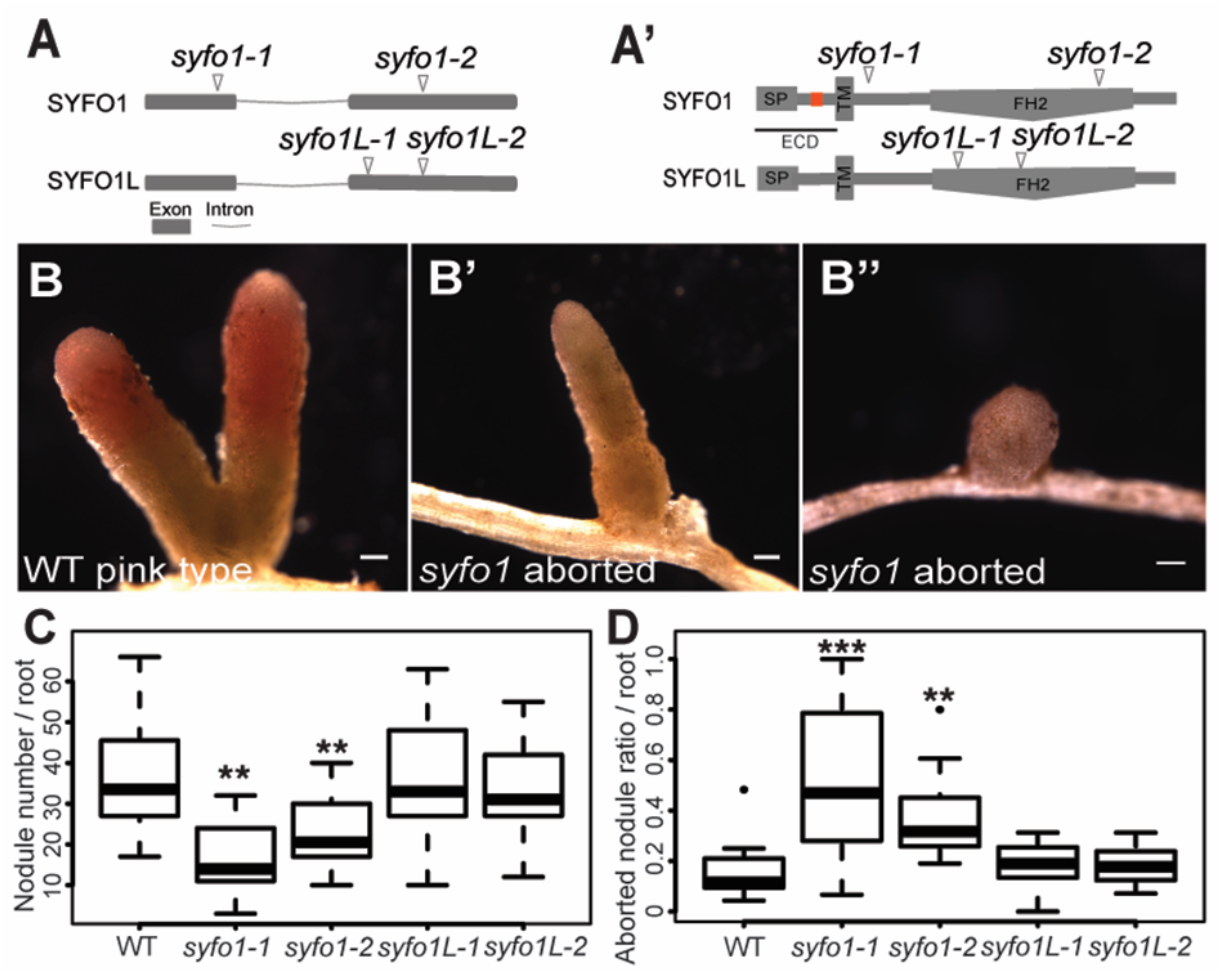
SYFO1 but not SYFO1L is required for nodulation. (*A-A’*) Isolation of independent mutant alleles with *Tnt1* transposon insertions mapping to different regions of the *SYFO1* and *SYFO1L* gene (*A*) and protein (*A’*) models, the extracellular domain is indicated as ECD and the orange box indicated the location of proline-rich repeat (PRR). (*B-B’’*) Nodule phenotypes observed on the different genotypes at 3 wpi with *S. meliloti* in open pots using WT R108 plants as a control. Scale bars indicate 200 μm. (*C-D*) Quantification of nodule numbers (*C*) and the ratio of aborted/wild-type like nodules (*D*) (n=10). Asterisks indicate a significant statistical difference based on a Tukey–Kramer multiple-comparison test with p-values <0.01 (**), < 0.001 (***). Data are shown as mean ± SE.

Although cytoskeleton-related mutants, like Lotus *pir1, nap1* and *scarn [34, 35]*, exhibit an impaired root hair growth, this was not observed for *syfo1-1* and *syfo1-2* where root hair length was indistinguishable from wild-type (Fig. S5). In agreement with this, we did not observe any differences in actin arrangement in growing root hairs within the symbiotically susceptible infection zone [4] under non-inoculated conditions (Fig. S6). These data demonstrate that SYFO1 is not required for normal root hair development under non-symbiotic conditions. As *SYFO1* transcripts were upregulated at 1-3 dpi, a stage where we observe most root hairs responding to the presence of the symbiont by root hair deformation (1-2 dpi) (Fig. 2A) and curling (2-3 dpi) (Fig. 2B), we phenotypically assessed *syfo1* mutants at these stages. Both mutant alleles showed significantly fewer deformations (Fig. 2C) as well as infection chambers and infection threads (Fig. 2E) at the respective time points. Both phenotypes were fully rescued when generating independent transgenic roots expressing a *ProSYFO1:SYFO1-GFP* (e.g. *syfo1-1c*) construct in these mutant backgrounds (Fig. 2D, F), demonstrating that the insertion in the *SYFO1* locus is causative for these phenotypes.

**Figure 2.**
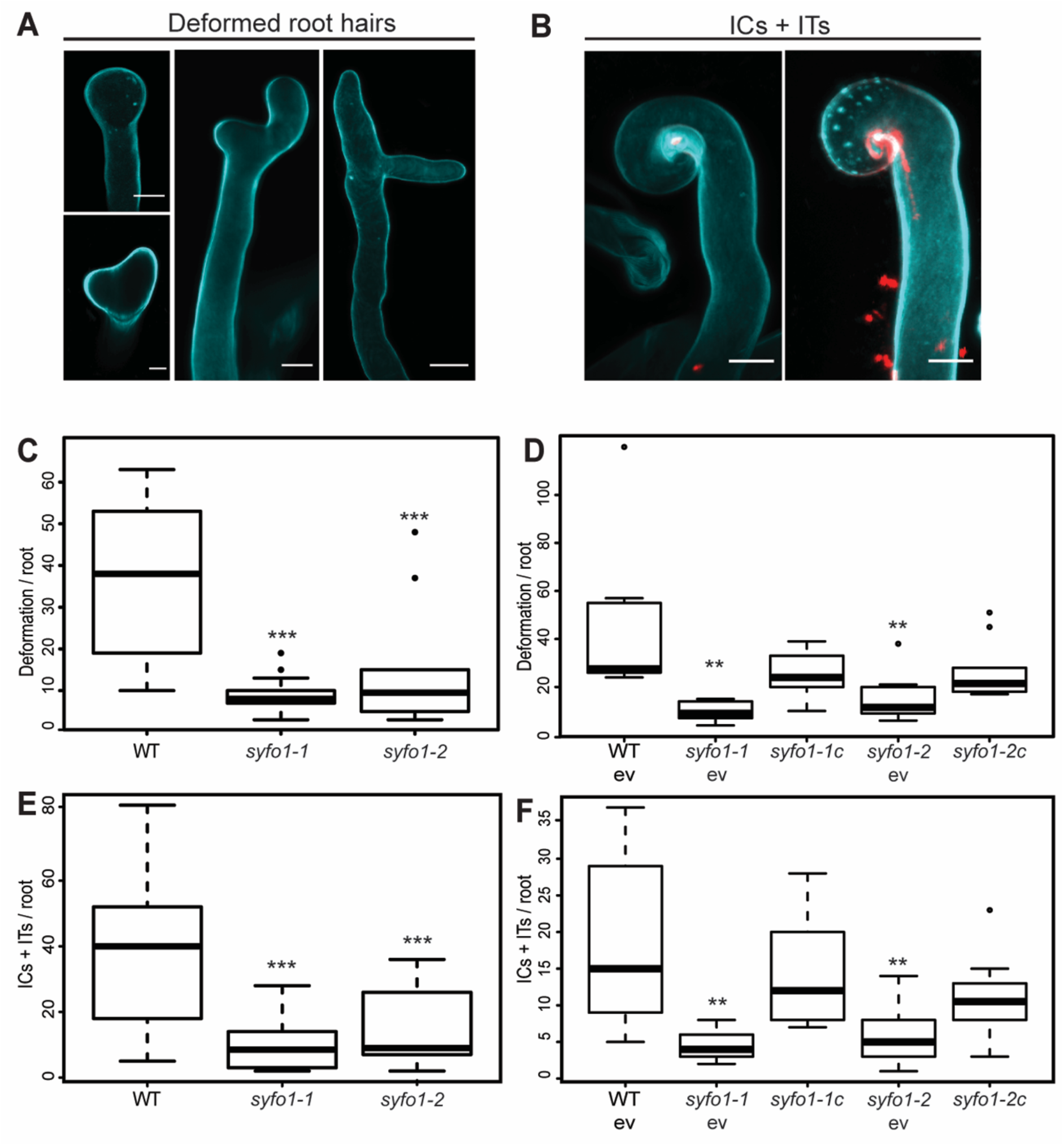
*syfo1* mutants are impaired in symbiotic root hair responses. Images show deformed root hairs (*A*), infection chambers (ICs) and infection threads (ITs) (*B*) on wild-type plants to illustrate the scored structures. Scale bars indicate 10 μm. *syfo1* mutants show significantly reduced responsiveness to the presence of compatible rhizobia when assessing root hair deformations (*C*) and infection-related structures such as ICs + ITs (*E*). These phenotypes were complemented by introducing a genomic version of a full-length *SYFO1* gene driven by the endogenous *SYFO1* promoter (*syfo1-1c*, *syfo1-2c*) with an empty vector (ev) transformation control aside (*D and F*). Asterisks indicate a significant statistical difference based on a Tukey–Kramer multiple-comparison test with p-values <0.01 (**), < 0.001 (***). Data are shown as mean ± SE with independent 9-14 plants for phenotypical analysis, and 10 plants for complementation analysis.

As it was previously demonstrated that the inoculation of Lotus root hairs with its symbiont *M. loti* resulted in a strong polarization and bundling of actin filaments with a strong accumulation of F-actin in the root hair tip [34, 35], we tested whether this pattern is affected in *syfo1* mutants. In the absence of *S. meliloti*, longitudinal actin filaments were observed in young growing root hairs within the infection zone of Medicago wild-type plants (Fig. S7A). In agreement with the observations in Lotus, actin strongly bundled and polarized with an accumulation of F-actin at the apex (Fig. S7B-C) or occasionally at the apical shank of responding root hairs (Fig. S7D) in almost 90% of all root systems of wild-type plants (Fig. S7E). However, this pattern was strongly reduced in both *syfo1* mutant alleles where only about 30% of all tested roots contained root hairs responding with the above-mentioned pattern (Fig. S7E).

### SYFO1 associates with polar actin assemblies under symbiotic conditions

Since our data indicated a symbiosis-specific role of SYFO1 in root hair polarization, we investigated spatial and temporal dynamics of SYFO1 at subcellular resolution. In roots expressing the *ProSYFO1:SYFO1-GFP* construct, we observed a weak homogenous signal of the SYFO1 protein at the PM of root hairs in the absence of rhizobia (Fig. 3A). The underlying low, basal expression was also detected by qRT-PCR (Fig. S3A) while it was most likely too weak when using the nuclear-localized GFP reporter to test promoter activity (Fig. S1). Interestingly, SYFO1 strongly accumulated in subapical and apical foci at root hair tips prior to deformation at 2 dpi with *S. meliloti* (Fig. 3B, C), which strongly resembled actin patterns observed upon Nod Factor application as reported earlier [34]. In root hairs that morphologically responded by deformation (Fig. 3D) and curling (Fig. 3E), SYFO1 distributed again along the PM with only mild accumulations at the apical region (Fig. 3D). SYFO1 also resided along the infection thread membrane even though to a much weaker extent (Fig. 3F-F’). When co-localizing actin and SYFO1 (here: *ProUbi-SYFO1-GFP*) we frequently observed transient SYFO1 accumulations in close proximity to enlarged nuclei, a hallmark for symbiotically activated root hairs [40], with actin bundles orienting towards a nucleation centre at the apical shank of the root hair (Fig. 3G-G’).

**Figure 3.**
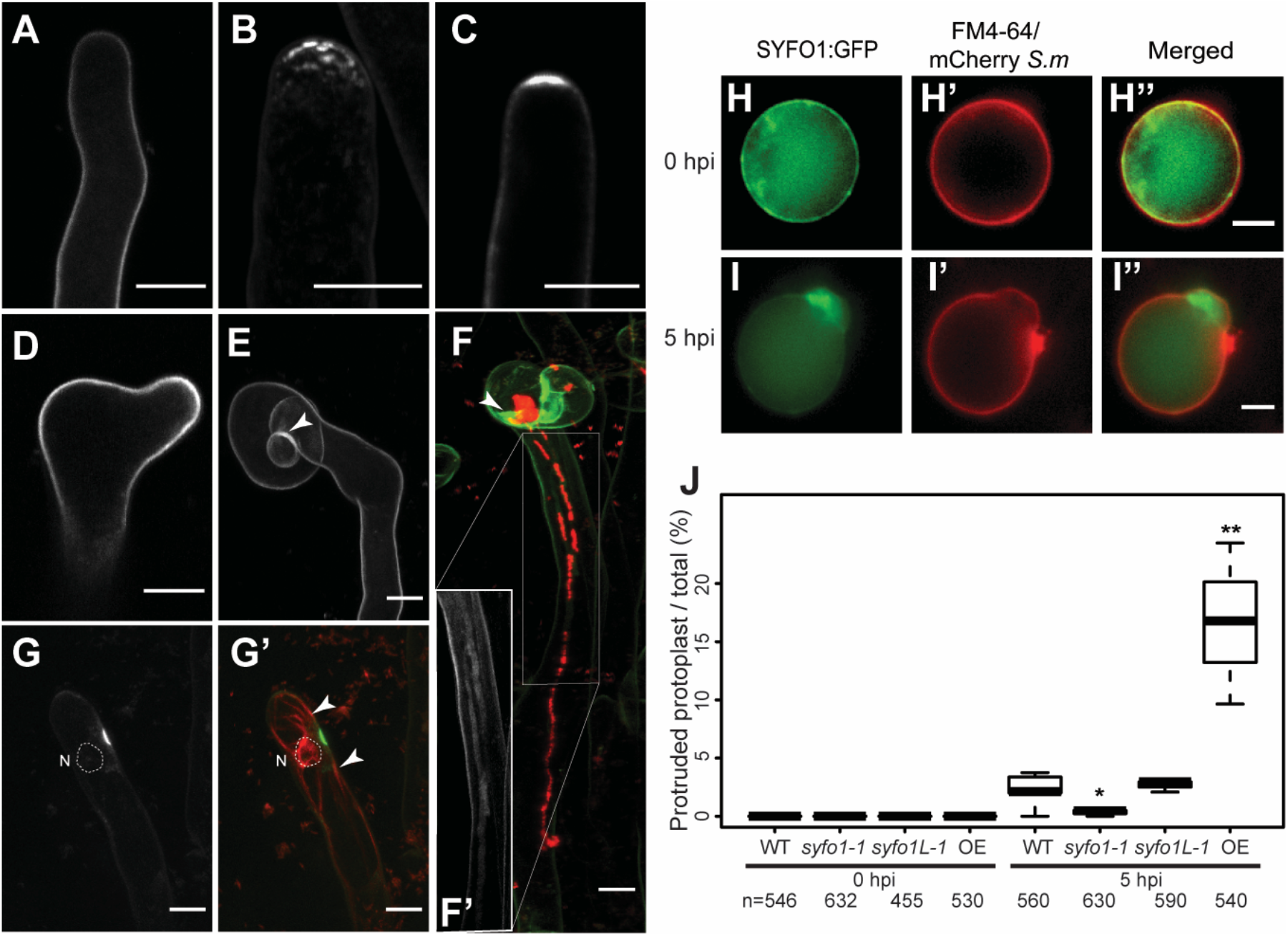
SYFO1 functions as a symbiotic polarity factor in root hairs. SYFO1-GFP localizes homogeneously to the PM of root hairs under mock conditions (*A*). At 2 dpi with *S. meliloti*, SYFO1 transiently polarizes at subapical (*B*) and apical regions (*C*) of root hairs, before distributing equally along the PM during root hair deformation (*D*) and curling (*E*). The arrowhead marks cell wall autofluorescence around the IC (*E and F*). SYFO1 remains on the IT membrane (*F* and inset *F’*). Co-localisation between SYFO1-GFP and actin marker ABD2:mCherry (*G-G’*). The nucleus encircled with a dash line in *G-G’* is based on corresponding transmitted light image (not shown). The arrowheads point towards actin bundles orienting towards a nucleation centre at the apical shank of the root hair. Scale bars indicate 10 μm. Protoplasts of a ROC expressing SYFO1 and counterstained with the styryl dye FM4-64 remain uniformly round at 0 dpi (*H-H’’*) while focal membrane protrusions with accumulated SYFO1 were found at 5 hpi with Ds-Red expressing *S. meliloti* (*I-I’’*). Scale bars indicate 5 μm. (*J*) Quantification of membrane protrusions in protoplasts using different genetic backgrounds. Asterisks indicate a significant statistical difference based on an ANOVA followed by a Fisher LSD test, with p-values <0.05 (*), < 0.01 (**). Data are shown as mean ± SE of 3 independent biological replicates with *n* indicating the total number of protoplasts being scored.

### Ligand-dependent morphological change mediated by SYFO1

To test whether SYFO1 acts in membrane-associated actin nucleation and polar filament elongation, we made use of the fact that extension of actin filaments can drive membrane protrusions as shown in cells lacking a rigid cell wall such as *Drosophila* Schneider 2 and mouse P19 cells [41, 42]. For this, we generated a transgenic Medicago root organ culture (ROC) constitutively expressing SYFO1-GFP for 8 weeks. When isolating protoplasts from this culture, SYFO1 resided in the PM where it co-localized with the membrane stain FM4-64 with some accumulations in the sub-membrane space (Fig. 3H-H’’). While no membrane protrusions were formed in the absence of bacteria, we observed these membrane outgrowths upon inoculation of protoplasts with *S. meliloti* for 5 hours. They coincided with SYFO1 accumulations inside these structures and often formed adjacent to bacterial aggregates (Fig. 3I-I’’). To unambiguously verify that SYFO1 is the key driver of these protrusions we isolated protoplasts from our *syfo1-1* and *syfo1L-1* mutants. No protrusions were found in the absence of rhizobia in any of the used genotypes. Upon inoculation of protoplasts with *S. meliloti*, those expressing wild-type *SYFO1* (wild-type, *syfo1L-1*) or over-expressing SYFO1-GFP (OE) developed protrusions while they were entirely absent on protoplasts isolated from the *syfo1-1* mutant (Fig. 3J). This demonstrates that focal accumulation of SYFO1 can drive cell polarisation and membrane deformations in a stimulus-dependent manner.

### A SYFO1-mediated cell wall-plasma membrane-cytoskeleton continuum is required for symbiotic responses in root hairs

As actin binding of formins is generally mediated by the FH2 domain that is also present in the cytosolic domain of SYFO1 (Fig. 1A’), we examined the extracellular region, which is less prominently found among formin proteins. Further sequence analysis of the SYFO1^ECD^ (extracellular domain) revealed the presence of a proline-rich repeat (PRR) (e.g. Ser-Pro-Pro-Pro-Ser-Pro-Ser-Ser [SPPPSPSS]) between SP and TMD, which resembles a canonical motif of extensin proteins that have been proposed to contribute to cell wall architecture and tensile strength [43, 44]. Thus we investigated a possible role in cell wall association of the ECD. For this, we fused the 82 amino acids of the SYFO1 ECD to GFP (SYFO1^ECD^-GFP) and expressed it in *Nicotiana benthamiana* leaf epidermal cells. The fluorescent label was found in the cell periphery in control cells (Fig. 4A-A’’) while it remained predominantly associated with the cell wall upon plasmolysis (Fig. 4B-B’’). In order to test whether this extensin-like motif contributes to the lateral immobilization of SYFO1 via cell wall anchoring we performed Fluorescence Recovery After Photobleaching (FRAP) experiments on root hairs constitutively expressing a full-length SYFO1 (SYFO1-GFP) or a mutant variant where we deleted the proline-rich repeat (SYFO1^ΔPRR^-GFP). These experiments revealed a slow recovery of the bleached region and a mobile fraction of about 24% for wild-type SYFO1 whereas the mobile fraction for SYFO1^ΔPRR^ was significantly higher (57%) (Fig. 4C-E). This clearly indicates that the PRR segment within the SYFO1 ECD anchors this formin to the cell wall. To address whether the cell wall association is required for SYFO1 function we conducted genetic complementation experiments, we generated transgenic roots expressing SYFO1^ΔPRR^ under the control of the native *SYFO1* promoter in our *syfo1-1* and *syfo1-2* alleles. In contrast to the full-length SYFO1 (Fig. 2), the deletion of the PRR fully abolished the ability to complement the root hair deformation phenotype of *syfo1* mutants (Fig. 4F), demonstrating that a SYFO1-mediated cell wall-plasma membrane-actin continuum is required for symbiotic responsiveness of root hairs in *M. truncatula*.

**Figure 4.**
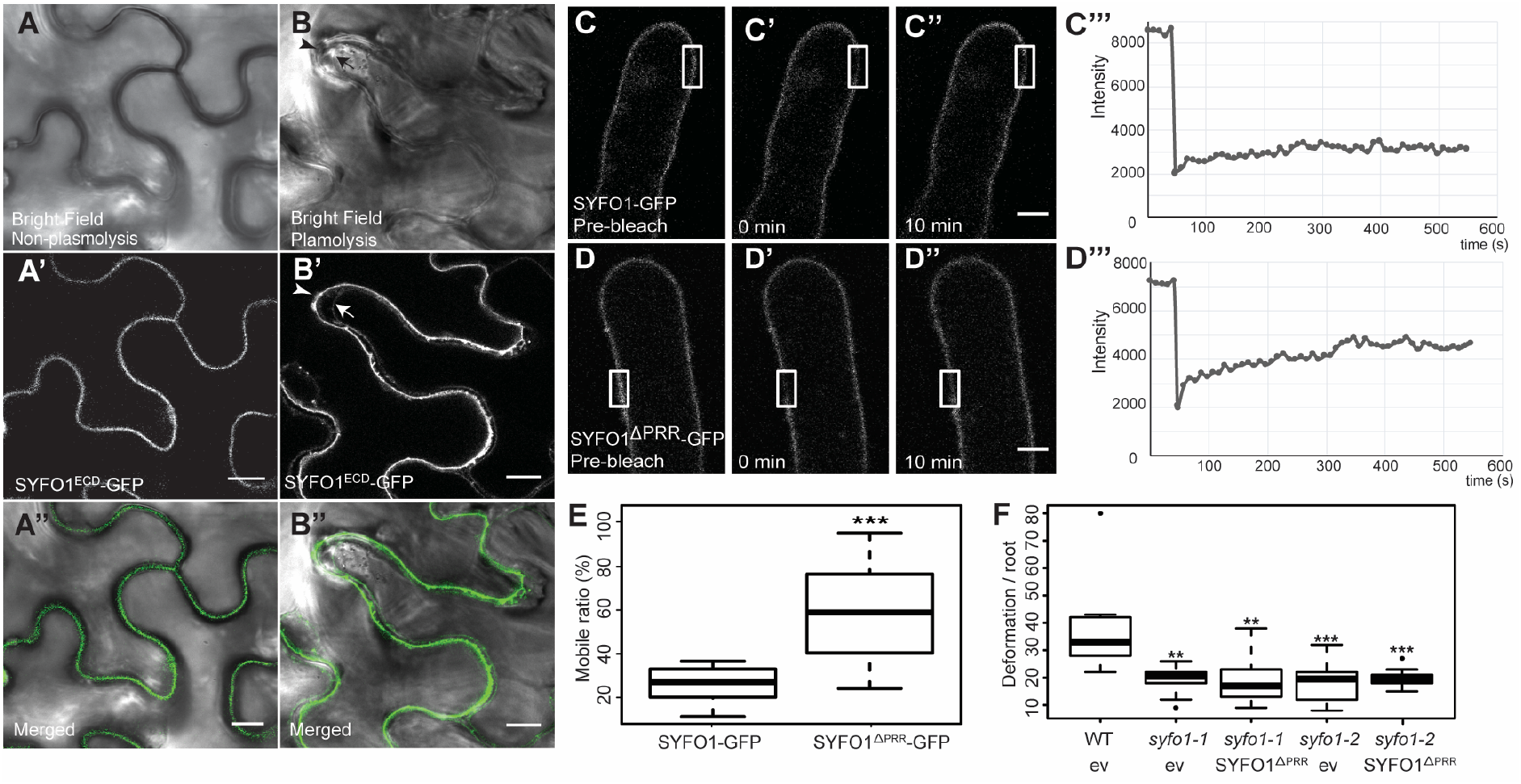
Cell wall association of SYFO1 is essential for its function. The constitutively expressed ECD of SYFO1 labelled the cell periphery in non-plasmolysed cells (*A-A’’*) and remained at the cell wall upon plasmolysis (*B-B’’*). Arrowheads and arrows mark the cell wall and the retracted plasma membrane, respectively. FRAP experiments on roots hairs revealed a low mobility of full-length SYFO1 (*C-C’’’*) while deletion of the PRR resulted in an increased mobility of the protein (*D-D’’’*). Scale bars indicate 10 μm. (*E*) Quantification of the mobile fractions of SYFO1 (n=17) and SYFO1^ΔPRR^ (n=12), asterisks indicate a significant statistical difference based on a student t test. (*F*) The SYFO1^ΔPRR^ variant failed to genetically complement both *syfo1* mutant alleles in comparison to roots transformed with the empty vector (ev) scoring n=10 independent root systems per genotype. Asterisks indicate a significant statistical difference based on a Tukey–Kramer multiple-comparison test with p-values <0.001 (***), p<0.01 (**). Data are shown as mean ± SE.

## DISCUSSION

The engulfment of a single rhizobium by a tightly curled root hair represents a fascinating process that allows most legumes to keep full control over the onset of root hair infection and might minimize the risk of infection by non-symbiotic bacteria. Considering that the formin protein SYFO1 serves symbiotic but not general functions in root hair responsiveness (Fig. S5 and Fig. S6) demonstrates that Medicago genetically re-wired few components of the actin machinery to drive root hair curling. However, the generic set of actin and polarity factors such as NAP1/PIR1, members of the SCAR/WAVE complex [34], and RHO OF PLANTS 10 [45] are additionally required for root hair growth *per se*. Consequently, these mutants show impaired growth also under non-symbiotic conditions. In contrast, legumes specifically evolved SCARN, another member of the SCAR/WAVE complex, which is required for rhizobial infection but not for growth under fully fertilized conditions [35]. The fact that the number of infection chambers was higher in Lotus *scarn* mutants compared to wild-type plants [35], places SCARN downstream of SYFO1. Since the bacterial colonization phenotypes in nodules are comparable between both mutants, both may contribute to actin function at this stage.

SYFO1 with its transmembrane helix and the extracellular proline-rich repeat carries all features of a group 1 formin [46]. As previously described for AtFH1 [47], the PRR mediates cell wall association and consequently restricts the lateral mobility also in SYFO1 (Fig. 4E). Its ability to initiate membrane protrusions in cell wall-depleted protoplasts further suggests that SYFO1 is involved in actin nucleation and filament elongation (Fig. 3G-G’; Fig. S8). In line with non-symbiotic formins, such as AtFH4 from Arabidopsis that re-localizes to infection sites of the powdery mildew fungus *Blumeria graminis* [48], we hypothesize that SYFO1 evolved to specifically mediate targeted secretion of cell wall constituents and other cargo material to sustain symbiotic root hair responses including root hair curling and later stages of infection. This entirely depends on the ability of SYFO1 to associate with the cell wall (Fig. 4) where it maintains a cell wall-plasma membrane-cytoskeleton continuum that cannot be functionally complemented by other, non-symbiotic formins that remain being expressed in *syfo1* mutants.

## MATERIALS AND METHODS

### Plant growths and phenotypical analysis

For phenotypical analysis *Medicago truncatula* wild-type R108, *syfo1-1*, *syfo1-2*, *syfo1L-1* and *syfo1L-2* seeds were scarified and sterilized before being sown on 1% agar plates for germination and kept in darkness at 4°C for 3 days for vernalization. Germination was allowed for up to 24 hours at 24°C before transferring the seedlings to plates containing Fahraeus medium [49] for 4 days in the presence of 1 mM nitrate before being transferred to a plate culture system without nitrogen for phenotyping studies. Plants were inoculated with 1ml *Sinorhizobium meliloti* 2011 (mCherry) at an OD_600_ of 0.05 (on plates or open pots with 1:1 ratio of vermiculite and sand mixture). Symbiotic responses including root hair deformations, infection chamber formation and IT development were scored 5 dpi of plants with *S. meliloti* on plates. Soil-based nodulation phenotyping samples were harvested and quantified at 3 wpi. Nodules were embedded in 7% Agar and sectioned with a thickness of 60 μm using a vibratome.

### Genotyping of *Tnt1* insertion lines and quantitative Real-Time PCR

R0 or R1 seeds of *M. truncatula* R108 *Tnt1* transposon insertion lines were obtained from the Noble Research Institute (OK, USA) and insertions were verified using primers listed in Table S4. Total RNA of control and insertion lines was extracted using a commercial kit (Spectrum™ Plant Total RNA Kit, Sigma life science) following the supplier’s instructions. An additional DNaseI treatment was performed. Synthesis of cDNA and qRT-PCR were conducted as described earlier [50] using the SuperScriptIII reverse Transcriptase (Invitrogen). All data were normalized to Ct values of the housekeeping gene ubiquitin [51] using primers listed in Table S4.

### Hairy root transformation and inoculation of rhizobia

*M. truncatula* hairy root transformation was performed as previously described [52] using the *Agrobacterium rhizogenes* strain ARqua1. Plants were transferred weekly to fresh plates containing Fahraeus medium (pH 6.0) supplemented with 0.5 mM NH_4_NO_3_ and followed by 2 days of growth on nitrogen-free Fahraeus medium containing 0.1 μM AVG prior to inoculation. Images for localisation studies and root hair phenotyping analyses were taken on plants inoculated for 2 days and 5 days, respectively.

### Phylogenetic and selective pressure analysis

SYFO1 (Medtr5g036540.1) and SYFO1L (Medtr8g062830.1) protein sequences were used as queries for a tBLASTn v2.8.1+ (10.1186/1471-2105-10-421) search against a database of 101 Angiosperms genomes (Table S2, sequences can be downloaded from the SymDB database (10.1038/s41477-020-0613-7): http://www.polebio.lrsv.ups-tlse.fr/symdb) with an e-value threshold of 1e-10. Sequences were then aligned using MAFFT v7.407 (10.1093/molbev/mst010) with default parameters. The resulting alignment was trimmed using trimAl v1.4 rev22 (10.1093/bioinformatics/btp348) to remove positions containing more than 20% of gaps. The cleaned alignment was then subjected to a Maximum Likelihood (ML) analysis using IQ-TREE v1.6.7 (10.1093/molbev/msu300) as described here after. First, the best-fitting evolutionary model was tested using ModelFinder (10.1038/nmeth.4285). Then a ML search was performed using 10,000 replicates of SH-aLRT (10.1093/sysbio/syq010) for testing branches support. The tree was finally visualized and annotated with iTOL v4.4 (10.1093/nar/gkw290).

Signal peptide and transmembrane domains were predicted from proteins using signal v5.0 (10.1038/s41587-019-0036-z) and TMHMM v2.0c (PMID: 9783223G) respectively using default parameters.

To look for relaxation (K<1) or intensification (K>1) of selection acting on different lineages of interest in Eudicots (Table S3), we used the RELAX program (10.1093/molbev/msu400). This method calculates different synonymous and non-synonymous substitution rates (ω = *d*N/*d*S) using the phylogenetic tree topology for both foreground and background branches. Because (i) the programs used does not accept gaps in codon sequences and (ii) there is a negative correlation between the number of sequences and the number of ungapped positions, we used different numbers of input sequences for RELAX analysis. Protein sequences from SYFO1 and SYFO1L orthologs were aligned using MUSCLE v3.8.382. Short sequences were excluded to maximize sequence number while limiting gapped positions compared to SYFO1 and SYFO1L sequences of Medicago using a custom R script. We opted for 151 CDS sequences corresponding to 1008 positions (Table S3).

### Construct design

The constructs used in this study were designed using Golden Gate cloning [53]. 2.5 kb upstream of the SYFO1 start codon were chosen as putative promoter region. A Golden Gate compatible full-length genomic DNA version (Medtr5g036540.1) was synthesized (GENEWIZ, Germany) by removing the *BpiI* and *BsaI* restriction sites via silent mutations. All cloning primers are listed in Table S4. To select transgenic roots a *pUbi:NLS-mCherry* or *pUbi:NLS-2xCerulean* cassette was additionally inserted into the different T-DNAs containing the transgenes of choice as previously described [50]. Level II and level III constructs were assembled based on the principle described earlier [53]. An overview about all designed constructs is provided in Table S5.

### Confocal Laser-Scanning Microscopy and FRAP

For imaging the NLS-GFP reporter module, sectioned nodules, protein localisation and plasmolysis we used a Leica TCS SP8 confocal microscope equipped with a 20x HCX PL APO water immersion lenses (Leica Microsystems, Mannheim, Germany). GFP was excited with a White Light Laser (WLL) at 488 nm and the emission was detected at 500-550 nm. mCherry fluorescence was excited using a WLL at 561nm and emission was detected between 575-630 nm. Samples, co-expressing two fluorophores were imaged in sequential mode between frames. FRAP analysis was conducted using a Zeiss LSM 880 Airyscan confocal microscope. For this, the VP (Virtual Pinhole) mode was adapted based on the fluorescence intensity of probe and Airyscan processing was performed using the ZEN (black edition) software package. The bleaching region, reference region and background region were selected at identical size. A pre-bleaching time of 5 seconds was chosen. Bleaching was set to stop upon the intensity dropping to 50% of the initial intensity before fluorescence recovery was recorded for 10 minutes. Using the FRAP data process package in ZEN (black version), the mobile fraction was calculated by the following equation (mobile fraction= *I*1/ *IE*), where *I*1 represents the dropped intensity and the *IE* represents the recovered intensity normalized to the reference intensity.

### Root organ culture, protoplast extraction, and inoculation

Transgenic Root Organ Culture (ROC) of *M. truncatula* expressing SYFO1-GFP were obtained via hairy root transformation according to [52]. Fully transformed root segments were cut and initially transferred to M-Media plates (Becard & Fortin, 1988) containing Augmentin (Sigma). 1 g of amoxicillin–200 mg of clavulanic acid) (400 mg/L) for two weeks and then subcultured on plates supplemented with 200 mg/L Augmentin for additional two weeks to remove *A. rhizogenes* contamination. Plates were sealed with micropore tape and incubated at 24°C in dark. Afterwards, the ROC was transferred to M-media plates without Augmentin to support faster tissue growth. Continuous expression of the transformation marker was monitored throughout the entire experiment.

Protoplasts were isolated from ROC by cutting the roots in small pieces of 2-5 mm length and processed as described previously [54]. For inoculation, an *S. meliloti* culture was diluted in W5 solution [54] to an OD_600_ of 0.05 before being added to the protoplasts.

### Actin phalloidin staining and plasma membrane FM4-64 staining

Phalloidin-based actin staining was performed according to a published protocol [31]. In brief, *Medicago* roots were transferred into Fahraeus medium containing 300 μM MBS (m-maleimidobenzoyl-N-hydroxysuccinimide ester) for 30 minutes to stabilize the actin filaments. The material was then fixed in 2 % formaldehyde in actin-stabilizing buffer (ASB) solution [34]. Phalloidin was added to a final concentration of 16 μM and staining was performed in the dark for 30 minutes. Root-derived protoplasts were submerged in a FM4-64 solution with a final concentration of 20 μM and incubated on ice for 5-10 mins before imaging.

## ACKNOWLEDGEMENTS

This study was supported by the project Engineering Nitrogen Symbiosis for Africa (ENSA) currently supported through a grant to the University of Cambridge by the Bill & Melinda Gates Foundation (OPP1172165) and UK government’s Department for International Development (DFID) as well as by the China Scholarship Council (CSC) (grant no. 201506350004 to PL). JK, CL and PMD belong to the LRSV, which is part of the TULIP LABEX (ANR-10-LABX-41). We thank the staff of the Life Imaging Center (LIC) in the Center for Biological Systems Analysis (ZBSA) of the Albert-Ludwigs-University of Freiburg for help with their confocal microscopy resources, and the excellent support in image recording. The *Medicago truncatula* plants utilized in this research project, which are jointly owned by the Centre National De La Recherche Scientifique, were obtained from Noble Research Institute, LLC and were created through research funded, in part, by a grant from the National Science Foundation, NSF-0703285. We are grateful to the genotoul bioinformatics platform Toulouse Occitanie (Bioinfo Genotoul, doi: 10.15454/1.5572369328961167E12) for providing computing resources.

We would explicitly thank all the members of our team for the fruitful discussions and for providing their individual expertise throughout the course of the project.

The authors declare no competing interests.

## Author Contributions

Conceptualization, P.L. and T.O.; Investigation, P.L., C.S., B.L., F.A.D., J.K., C.L., P.M.D. and T.O.; Writing –Original Draft, P.L. and T.O.; Writing –Review & Editing, P.L., C.S., B.L., F.A.D., J.K., C.L., P.M.D. and T.O.; Funding Acquisition, P.L., P.M.D. and T.O.; Supervision, T.O.

## REFERENCES

1. Mylona, P., Pawlowski, K., and Bisseling, T. (1995). Symbiotic Nitrogen-Fixation. Plant Cell 7, 869–885.

2. Oldroyd, G.E.D., and Downie, J.M. (2008). Coordinating nodule morphogenesis with rhizobial infection in legumes. Annu Rev Plant Biol 59, 519–546.

3. Brewin, N.J. (2004). Plant cell wall remodelling in the rhizobium-legume symbiosis. Crit Rev Plant Sci 23, 293–316.

4. Gage, D.J. (2004). Infection and invasion of roots by symbiotic, nitrogen-fixing rhizobia during nodulation of temperate legumes. Microbiol Mol Biol R 68, 280–+.

5. Esseling, J.J., Lhuissier, F.G.P., and Emons, A.M.C. (2003). Nod factor-induced root hair curling: Continuous polar growth towards the point of nod factor application. Plant Physiology 132, 1982–1988.

6. Oldroyd, G.E.D., and Downie, J.A. (2004). Calcium, kinases and nodulation signalling in legumes. Nat Rev Mol Cell Bio 5, 566–576.

7. Bibikova, T.N., Blancaflor, E.B., and Gilroy, S. (1999). Microtubules regulate tip growth and orientation in root hairs of Arabidopsis thaliana. Plant Journal 17, 657–665.

8. Yang, G.H., Gao, P., Zhang, H., Huang, S.J., and Zheng, Z.L. (2007). A Mutation in MRH2 Kinesin Enhances the Root Hair Tip Growth Defect Caused by Constitutively Activated ROP2 Small GTPase in Arabidopsis. Plos One 2.

9. Yi, K., Guo, C., Chen, D., Zhao, B., Yang, B., and Ren, H. (2005). Cloning and functional characterization of a formin-like protein (AtFH8) from Arabidopsis. Plant Physiol 138, 1071–1082.

10. Jones, M.A., Shen, J.J., Fu, Y., Li, H., Yang, Z.B., and Grierson, C.S. (2002). The Arabidopsis Rop2 GTPase is a positive regulator of both root hair initiation and tip growth. Plant Cell 14, 763–776.

11. Parker, J.S., Cavell, A.C., Dolan, L., Roberts, K., and Grierson, C.S. (2000). Genetic interactions during root hair morphogenesis in Arabidopsis. Plant Cell 12, 1961–1974.

12. Carol, R.J., Takeda, S., Linstead, P., Durrant, M.C., Kakesova, H., Derbyshire, P., Drea, S., Zarsky, V., and Dolan, L. (2005). A RhoGDP dissociation inhibitor spatially regulates growth in root hair cells. Nature 438, 1013–1016.

13. Fournier, J., Teillet, A., Chabaud, M., Ivanov, S., Genre, A., Limpens, E., de Carvalho-Niebel, F., and Barker, D.G. (2015). Remodeling of the infection chamber before infection thread formation reveals a two-step mechanism for rhizobial entry into the host legume root hair. Plant Physiology 167, 1233–1242.

14. Xie, F., Murray, J.D., Kim, J., Heckmann, A.B., Edwards, A., Oldroyd, G.E.D., and Downie, A. (2012). Legume pectate lyase required for root infection by rhizobia. P Natl Acad Sci USA 109, 633–638.

15. Gage, D.J., and Margolin, W. (2000). Hanging by a thread: invasion of legume plants by rhizobia. Curr Opin Microbiol 3, 613–617.

16. Heckmann, A.B., Lombardo, F., Miwa, H., Perry, J.A., Bunnewell, S., Parniske, M., Wang, T.L., and Downie, J.A. (2006). Lotus japonicus nodulation requires two GRAS domain regulators, one of which is functionally conserved in a non-legume. Plant Physiology 142, 1739–1750.

17. Bek, A.S., Sauer, J., Thygesen, M.B., Duus, J.O., Petersen, B.O., Thirup, S., James, E., Jensen, K.J., Stougaard, J., and Radutoiu, S. (2010). Improved Characterization of Nod Factors and Genetically Based Variation in LysM Receptor Domains Identify Amino Acids Expendable for Nod Factor Recognition in Lotus spp. Molecular Plant-Microbe Interactions 23, 58–66.

18. Madsen, L.H., Tirichine, L., Jurkiewicz, A., Sullivan, J.T., Heckmann, A.B., Bek, A.S., Ronson, C.W., James, E.K., and Stougaard, J. (2010). The molecular network governing nodule organogenesis and infection in the model legume Lotus japonicus. Nat Commun 1, 10.

19. Cerri, M.R., Wang, Q.H., Stolz, P., Folgmann, J., Frances, L., Katzer, K., Li, X.L., Heckmann, A.B., Wang, T.L., Downie, J.A., et al. (2017). The ERN1 transcription factor gene is a target of the CCaMK/CYCLOPS complex and controls rhizobial infection in Lotus japonicus. New Phytologist 215, 323–337.

20. Kawaharada, Y., James, E.K., Kelly, S., Sandal, N., and Stougaard, J. (2017). The ethylene responsive factor required for nodulation 1 (ERN1) transcription factor is required for infection-thread formation in Lotus japonicus. Molecular plant-microbe interactions 30, 194–204.

21. Schauser, L., Roussis, A., Stiller, J., and Stougaard, J. (1999). A plant regulator controlling development of symbiotic root nodules. Nature 402, 191–195.

22. Combier, J.P., Frugier, F., de Billy, F., Boualem, A., El-Yahyaoui, F., Moreau, S., Vernie, T., Ott, T., Gamas, P., Crespi, M., et al. (2006). MtHAP2-1 is a key transcriptional regulator of symbiotic nodule development regulated by microRNA169 in Medicago truncatula. Gene Dev 20, 3084–3088.

23. Ben Amor, B. (2003). The NFP locus of Medicago truncatula controls an early step of Nod factor signal transduction upstream of a rapid calcium flux and root hair deformation (vol 34, pg 495, 2003). Plant Journal 35, 140–140.

24. Limpens, E., Franken, C., Smit, P., Willemse, J., Bisseling, T., and Geurts, R. (2003). LysM domain receptor kinases regulating rhizobial Nod factor-induced infection. Science 302, 630–633.

25. Madsen, E.B., Madsen, L.H., Radutoiu, S., Olbryt, M., Rakwalska, M., Szczyglowski, K., Sato, S., Kaneko, T., Tabata, S., Sandal, N., et al. (2003). A receptor kinase gene of the LysM type is involved in legume perception of rhizobial signals. Nature 425, 637–640.

26. Arrighi, J.F., Barre, A., Ben Amor, B., Bersoult, A., Soriano, L.C., Mirabella, R., de Carvalho-Niebel, F., Journet, E.P., Gherardi, M., Huguet, T., et al. (2007). The Medicago truncatula lysine motif-receptor-like kinase gene family includes NFP and new nodule-expressed genes (vol 142, pg 265, 2006). Plant Physiology 143, 1078–1078.

27. Smit, P., Limpens, E., Geurts, R., Fedorova, E., Dolgikh, E., Gough, C., and Bisseling, T. (2007). Medicago LYK3, an entry receptor in rhizobial nodulation factor signaling. Plant Physiology 145, 183–191.

28. Radutoiu, S., Madsen, L.H., Madsen, E.B., Felle, H.H., Umehara, Y., Gronlund, M., Sato, S., Nakamura, Y., Tabata, S., Sandal, N., et al. (2003). Plant recognition of symbiotic bacteria requires two LysM receptor-like kinases. Nature 425, 585–592.

29. de Ruijter, N.C.A., Rook, M.B., Bisseling, T., and Emons, A.M.C. (1998). Lipochito-oligosaccharides re-initiate root hair tip growth in Vicia sativa with high calcium and spectrin-like antigen at the tip. Plant Journal 13, 341–350.

30. Cardenas, L., Feijo, J.A., Kunkel, J.G., Sanchez, F., Holdaway-Clarke, T., Hepler, P.K., and Quinto, C. (1999). Rhizobium Nod factors induce increases in intracellular free calcium and extracellular calcium influxes in bean root hairs. Plant Journal 19, 347–352.

31. Miller, D.D., de Ruijter, N.C.A., Bisseling, T., and Emons, A.M.C. (1999). The role of actin in root hair morphogenesis: studies with lipochito-oligosaccharide as a growth stimulator and cytochalasin as an actin perturbing drug. Plant Journal 17, 141–154.

32. Miller, D.D., Klooster, H.B.L.-t., and Emons, A.M.C. (2000). Lipochito-oligosaccharide nodulation factors stimulate cytoplasmic polarity with longitudinal endoplasmic reticulum and vesicles at the tip in vetch root hairs. Molecular plant-microbe interactions 13, 1385–1390.

33. Zepeda, I., Sanchez-Lopez, R., Kunkel, J.G., Banuelos, L.A., Hernandez-Barrera, A., Sanchez, F., Quinto, C., and Cardenas, L. (2014). Visualization of Highly Dynamic F-Actin Plus Ends in Growing Phaseolus vulgaris Root Hair Cells and Their Responses to Rhizobium etli Nod Factors. Plant Cell Physiol 55, 580–592.

34. Yokota, K., Fukai, E., Madsen, L.H., Jurkiewicz, A., Rueda, P., Radutoiu, S., Held, M., Hossain, M.S., Szczyglowski, K., Morieri, G., et al. (2009). Rearrangement of Actin Cytoskeleton Mediates Invasion of Lotus japonicus Roots by Mesorhizobium loti. Plant Cell 21, 267–284.

35. Qiu, L.P., Lin, J.S., Xu, J., Sato, S., Parniske, M., Wang, T.L., Downie, J.A., and Xie, F. (2015). SCARN a Novel Class of SCAR Protein That Is Required for Root-Hair Infection during Legume Nodulation. Plos Genet 11.

36. Zepeda, I., Sanchez-Lopez, R., Kunkel, J.G., Banuelos, L.A., Hernandez-Barrera, A., Sanchez, F., Quinto, C., and Cardenas, L. (2014). Visualization of highly dynamic F-actin plus ends in growing phaseolus vulgaris root hair cells and their responses to Rhizobium etli nod factors. Plant Cell Physiol 55, 580–592.

37. Yang, C., Czech, L., Gerboth, S., Kojima, S.I., Scita, G., and Svitkina, T. (2007). Novel roles of formin mDia2 in lamellipodia and filopodia formation in motile cells. Plos Biology 5, 2624–2645.

38. Larrainzar, E., Riely, B.K., Kim, S.C., Carrasquilla-Garcia, N., Yu, H.J., Hwang, H.J., Oh, M., Kim, G.B., Surendrarao, A.K., Chasman, D., et al. (2015). Deep Sequencing of the Medicago truncatula Root Transcriptome Reveals a Massive and Early Interaction between Nodulation Factor and Ethylene Signals. Plant Physiology 169, 233–+.

39. van Velzen, R., Holmer, R., Bu, F.J., Rutten, L., van Zeijl, A., Liu, W., Santuari, L., Cao, Q.Q., Sharma, T., Shen, D.F., et al. (2018). Comparative genomics of the nonlegume Parasponia reveals insights into evolution of nitrogen-fixing rhizobium symbioses. P Natl Acad Sci USA 115, E4700–E4709.

40. Oldroyd, G.E.D. (2013). Speak, friend, and enter: signalling systems that promote beneficial symbiotic associations in plants. Nature Reviews Microbiology 11, 252–263.

41. Matusek, T., Gombos, R., Szecsenyi, A., Sanchez-Soriano, N., Czibula, A., Pataki, C., Gedai, A., Prokop, A., Rasko, I., and Mihaly, J. (2008). Formin proteins of the DAAM subfamily play a role during axon growth. J Neurosci 28, 13310–13319.

42. Gombos, R., Migh, E., Antal, O., Mukherjee, A., Jenny, A., and Mihaly, J. (2015). The Formin DAAM Functions as Molecular Effector of the Planar Cell Polarity Pathway during Axonal Development in Drosophila. J Neurosci 35, 10154–10167.

43. Estevez, J.M., Kieliszewski, M.J., Khitrov, N., and Somerville, C. (2006). Characterization of synthetic hydroxyproline-rich proteoglycans with arabinogalactan protein and extensin motifs in Arabidopsis. Plant Physiol 142, 458–470.

44. Lamport, D.T., Kieliszewski, M.J., Chen, Y., and Cannon, M.C. (2011). Role of the extensin superfamily in primary cell wall architecture. Plant Physiol 156, 11–19.

45. Lei, M.J., Wang, Q., Li, X.L., Chen, A.M., Luo, L., Xie, Y.J., Li, G., Luo, D., Mysore, K.S., Wen, J.Q., et al. (2015). The Small GTPase ROP10 of Medicago truncatula Is Required for Both Tip Growth of Root Hairs and Nod Factor-Induced Root Hair Deformation. Plant Cell 27, 806–822.

46. Deeks, M.J., Hussey, P.J., and Davies, B. (2002). Formins: intermediates in signal-transduction cascades that affect cytoskeletal reorganization. Trends in Plant Science 7, 492–498.

47. Martiniere, A., Gayral, P., Hawes, C., and Runions, J. (2011). Building bridges: formin1 of Arabidopsis forms a connection between the cell wall and the actin cytoskeleton. Plant Journal 66, 354–365.

48. Sassmann, S., Rodrigues, C., Milne, S.W., Nenninger, A., Allwood, E., Littlejohn, G.R., Talbot, N.J., Soeller, C., Davies, B., Hussey, P.J., et al. (2018). An Immune-Responsive Cytoskeletal-Plasma Membrane Feedback Loop in Plants. Curr Biol 28, 2136–2144 e2137.

49. Fahraeus, G. (1957). The Infection of Clover Root Hairs by Nodule Bacteria Studied by a Simple Glass Slide Technique. J Gen Microbiol 16, 374–&.

50. Liang, P., Stratil, T.F., Popp, C., Marin, M., Folgmann, J., Mysore, K.S., Wen, J., and Ott, T. (2018). Symbiotic root infections in Medicago truncatula require remorin-mediated receptor stabilization in membrane nanodomains. Proc Natl Acad Sci U S A 115, 5289–5294.

51. Satge, C., Moreau, S., Sallet, E., Lefort, G., Auriac, M.C., Rembliere, C., Cottret, L., Gallardo, K., Noirot, C., Jardinaud, M.F., et al. (2016). Reprogramming of DNA methylation is critical for nodule development in Medicago truncatula. Nature Plants 2.

52. Boisson-Dernier, A., Chabaud, M., Garcia, F., Becard, G., Rosenberg, C., and Barker, D.G. (2001). Agrobacterium rhizogenes-transformed roots of Medicago truncatula for the study of nitrogen-fixing and endomycorrhizal symbiotic associations. Mol Plant Microbe Interact 14, 695–700.

53. Binder, A., Lambert, J., Morbitzer, R., Popp, C., Ott, T., Lahaye, T., and Parniske, M. (2014). A Modular Plasmid Assembly Kit for Multigene Expression, Gene Silencing and Silencing Rescue in Plants. Plos One 9.

54. Jia, N., Zhu, Y.L., and Xie, F. (2018). An Efficient Protocol for Model Legume Root Protoplast Isolation and Transformation. Frontiers in Plant Science 9.

