## Supplemental Figures for "A formin-mediated cell wall- plasma membrane- cytoskeleton continuum is required for symbiotic infections in *Medicago truncatula*"

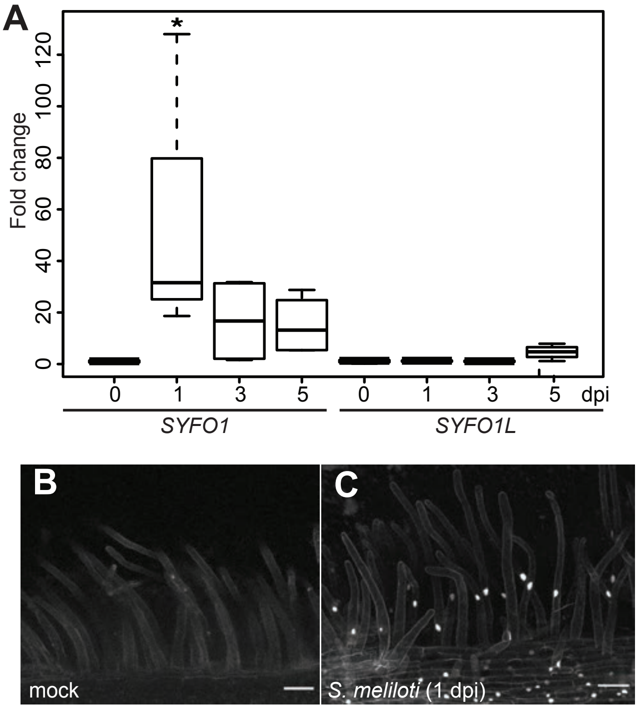


**Supplemental Figure S1. SYFO1 contributes to the symbiotic establishment.** (*A*) *SYFO1* and *SYFO1L* transcript levels before inoculation, at 1 dpi, 3 dpi and 5 dpi post inoculation. (*B-C*) Promoter activity was assessed with cellular resolution in transgenic *M. truncatula* root hairs by using the genetically encoded reporter *ProSYFO1:NLS-2xGFP*. Quantitative qRT-PCR was performed on cDNAs obtained from roots, with 4 biological replicates (each replicate contained roots from four independent plants). The graphs represent the fold change of ΔΔCt values obtained by qRT-PCR relative to ubiquitin. Asterisks indicate a significant statistical difference based on a Tukey–Kramer multiple-comparison test with p-values <0.05 (*). Scale bars indicate 50 µm (*B-C*).

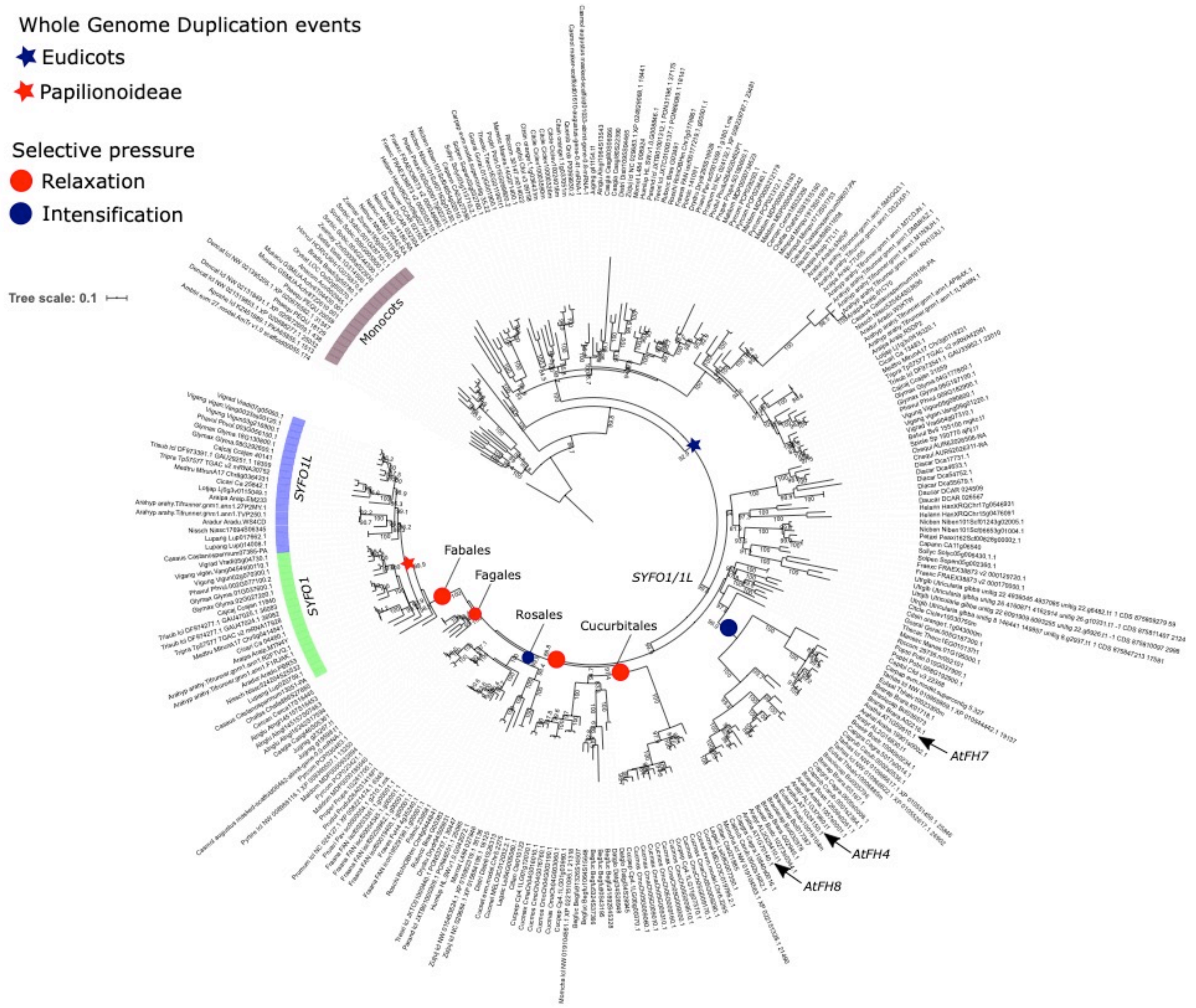


**Supplemental Figure S2. Maximum Likelihood tree (model: TIM3+F+R6; Log-likelihood: -124347.2322) of SYFO1 and SYFO1L family in 102 Angiosperm species.** The tree was rooted on *Amborella trichopoda*. The Eudicots duplication leading to the emergence of the SYFO1/SYFO1L clade is indicated with a blue star and the Papilionoideae duplication from which SYFO1 and SYFO1L derived is indicated with a red star. Monocots, SYFO1 and SYFO1L clades are indicated by violet, green and blue ribbons respectively. Results of the selective pressure analysis are marked as follow: red dots indicate relaxation (K<1 and p-val < 0.01) of the selection in the clade while blue dots stand for an intensification of the selection (K>1 and p-val < 0.01). Details of selective pressure analysis are presented in Table S3.

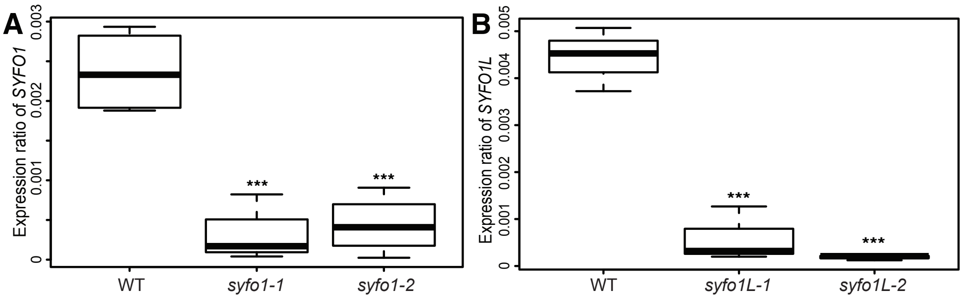


**Supplemental Figure S3. Determining transcript levels in Tnt1 insertion lines.** *SYFO1* transcript levels in *syfo1-1*, *syfo1-2*, *syfo1L-1*, *syfo1L-2* and wild-type R108 plants. Quantitative qRT-PCR was performed on cDNAs obtained from roots, with 4 biological replicates (each replicate contained roots from four independent plants). The graphs represent ΔΔCt values obtained by qRT-PCR relative to ubiquitin. Asterisks indicate a significant statistical difference based on a Tukey–Kramer multiple-comparison test with p-values <0.001 (***).

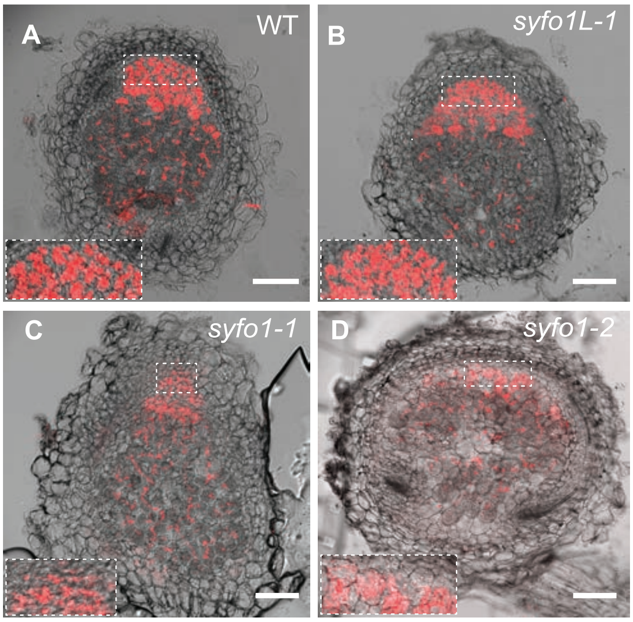


**Supplemental Figure S4. *syfo1* but not *syfo1L* mutants are impaired in early nodule development.** Semi-thin (60 µm) sections representative for 4 nodules that have been sectioned and embedded in low melting agarose from wild-type (*A*), *syfo1L* (*B*) and *syfo1* (*C-D)*. The dashed rectangles indicate the magnified selected zones. Plants were grown in open pots and inoculated for 3 weeks with *S. meliloti* (red) prior to the analysis. Scale bars indicate 100 µm.

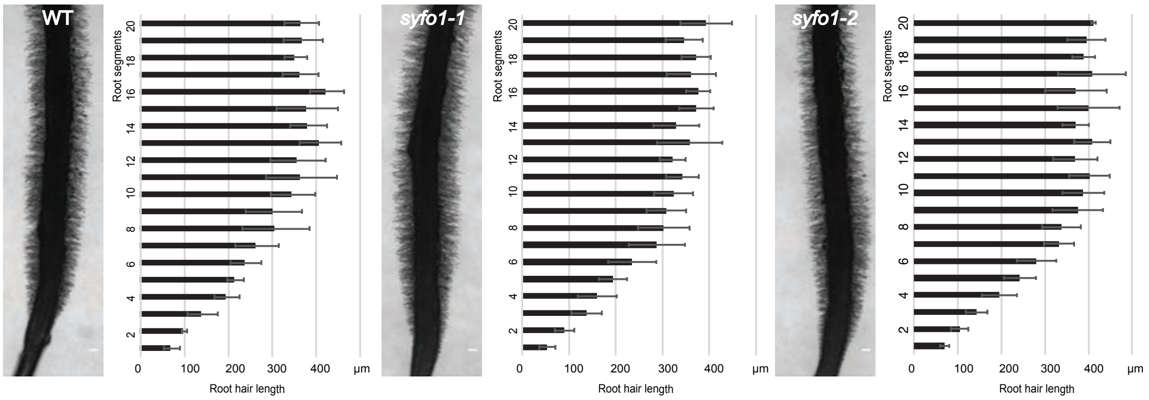


**Supplemental Figure S5. Root hairs develop normally on *syfo1-1* and *syfo1-2* mutants.** Plants were grown on vertical agar plates in the absence of rhizobia for 10 days prior to the analysis. Root hair length was measures in 20 consecutive root segments covering the designated infection zone (3 mm above the root hair tip). Scale bars indicate 200 µm. No statistically significant differences were found based on an ANOVA followed by a Fisher LSD test when comparing root hair length in between the corresponding root segment of the different genotypes. Data are shown as mean ± SE of 4 independent biological replicates with 4 root hairs being scored per experiment and genotype.

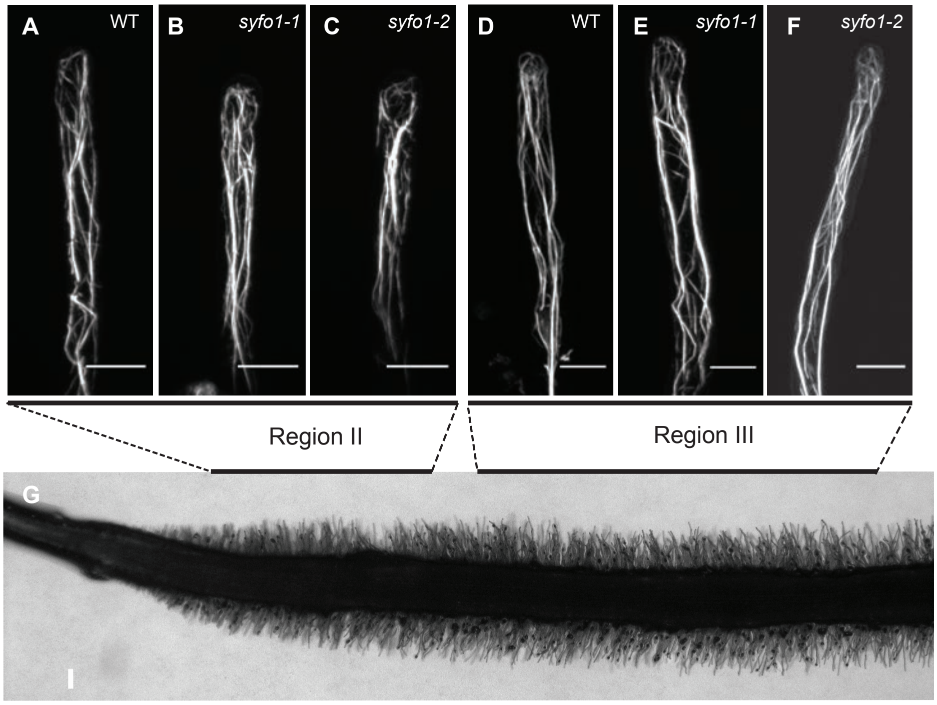


**Supplemental Figure S6. Actin arrangements are indistinguishable in young, uninoculated root hairs of wild type R108, *syfo1-1* and *syfo1-2* mutants.** Phalloidin staining of actin in root hairs of the differentiation zone (region II; *A-C*) and elongation zone (region III; *D-F*) as illustrated in *G*. Scale bars indicate 10 µm (*A-F*) and 200 µm (*G*).

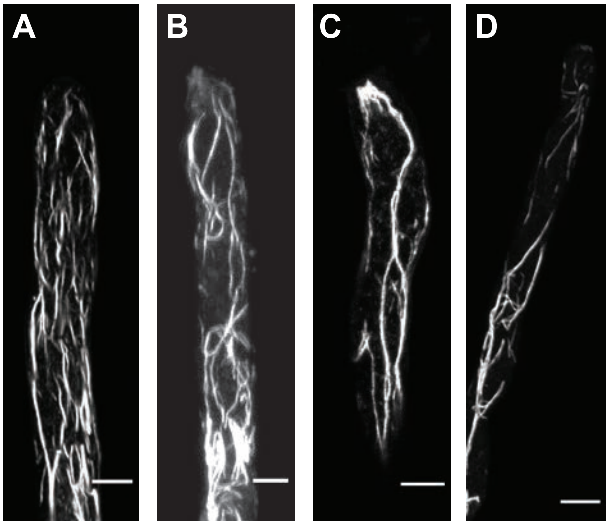


E

| genotype | $n_{polar}$ | $n_{total}$ | % of $n_{total}$ |
|---|---|---|---|
| WT | 35 | 40 | 88 |
| *syfo1-1* | 11 | 35 | 31 |
| *syfo1-2* | 5 | 20 | 25 |

**Supplemental Figure S7. Analysis of actin polarization upon rhizobial inoculation.** Actin patterns were visualized by Phalloidin staining of root hairs and observed under non-inoculated (*A*) and inoculated conditions after 2 dpi with *S. meliloti* (*B-D*). Representative images for longitudinal alignment of actin filaments in the absence of rhizobia in wild type (*A*), apically polarized actin (*B, C*) and polarized actin at the apical shank of root hairs (*D*) upon inoculation of wild-type root hairs. The appearance of polarized patterns ($n_{polar}$) as shown in panels *B-D* was scored on wild-type and *syfo1* mutant roots by absent/present calls of appearance on individual root systems. Scale bars indicate 10 µm (*A-D*).

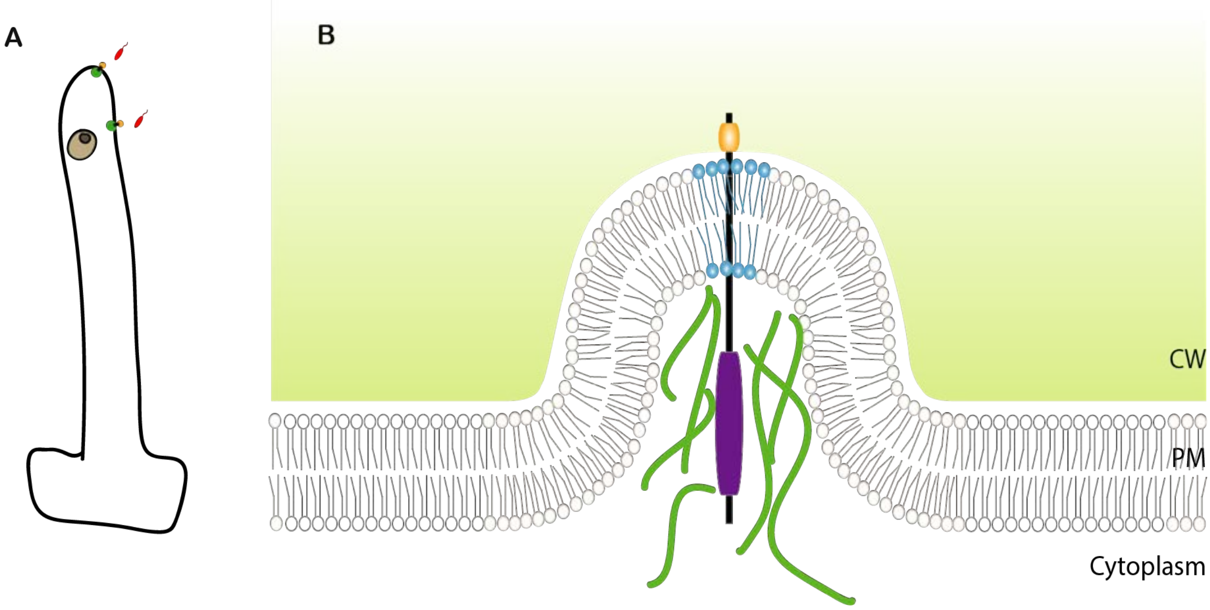


**Supplemental Figure S8. Proposed model for a SYFO1-mediated actin-plasma membrane-cell wall-continuum.** Our data suggest that SYFO1 establishes actin nucleation centres at the tip or the apical part of the shank of growing root hairs (green) upon inoculation with rhizobia (red) (*A*). (*B*) Association with the cell wall (CW; light green) by the proline-rich extracellular domain of SYFO1 (orange) laterally immobilizes SYFO1 and allows membrane deformations by SYFO1-mediated actin nucleation (green filaments) and formation of a polarity domain (bright blue) at these sites. PM= plasma membrane. The FH2 domain is depicted in purple.
